# Single-Cell Transcriptome Analysis of Lineage Diversity and Microenvironment in High-Grade Glioma

**DOI:** 10.1101/250704

**Authors:** Jinzhou Yuan, Hanna Mendes Levitin, Veronique Frattini, Erin C. Bush, Deborah M. Boyett, Jorge Samanamud, Michele Ceccarelli, Athanassios Dovas, George Zanazzi, Peter Canoll, Jeffrey N. Bruce, Anna Lasorella, Antonio Iavarone, Peter A. Sims

## Abstract

**Background:** Despite extensive molecular characterization, we lack a comprehensive understanding of lineage identity, differentiation, and proliferation in high-grade gliomas (HGGs). We sampled the cellular milieu of HGGs with massively-parallel single-cell RNA-Seq.

**Results:** While HGG cells can resemble glia or even immature neurons and form branched lineage structures, mesenchymal transformation results in unstructured populations. Glioma cells in a subset of mesenchymal tumors lose their neural lineage identity, express inflammatory genes, and co-exist with marked myeloid infiltration, reminiscent of molecular interactions between glioma and immune cells established in animal models. Additionally, we discovered a tight coupling between lineage resemblance and proliferation among malignantly transformed cells. Glioma cells that resemble oligodendrocyte progenitors, which proliferate in the brain, are often found in the cell cycle. Conversely, glioma cells that resemble astrocytes, neuroblasts, and oligodendrocytes, which are non-proliferative in the brain, are generally non-cycling in tumors.

**Conclusions:** These studies reveal a relationship between cellular identity and proliferation in HGG and distinct population structures that reflects the extent of neural and non-neural lineage resemblance among malignantly transformed cells.

## BACKGROUND

Gliomas are the most common malignant brain tumors in adults. High-grade gliomas (HGGs), which include Grade III anaplastic astrocytomas and Grade IV glioblastomas (GBMs), the deadliest form of brain tumor, are notoriously heterogeneous at the cellular level(1–5). While it is well-established that transformed cells in HGG resemble glia(6,7), the extent of neural lineage heterogeneity within individual tumors has not been thoroughly characterized. Furthermore, many studies have implied the existence of glioma stem cells – a rare subpopulation that is capable of self-renewal and giving rise to the remaining glioma cells in the tumor(8). Finally, the immune cells in the tumor microenvironment belong primarily to the myeloid lineage and drive tumor progression(9). However, little is known about the diversity of immune populations that infiltrate HGGs and a potential role of immune cells for immunotherapeutic approaches in HGG remains elusive(10).Therefore, questions about the nature and extent of interaction between transformed cells and the immune microenvironment in HGG persist despite extensive molecular profiling of bulk tumor specimens(3,7,11). Single-cell RNA-Seq (scRNA-Seq) approaches are shedding light on immune cell diversity in healthy contexts(12), and marker discovery for brain resident and glioma-infiltrating immune populations is an area of active study(13,14). Pioneering work used scRNA-Seq to provide a snapshot of the formidable heterogeneity characterizing human GBM(4,15,16). However, these early studies employed relatively low-throughput scRNA-Seq analysis which lacked the resolution necessary to deconvolve the full complexity of tumor and immune cells within individual HGGs. Later single cell studies in glioma focused on lower-grade gliomas and the effects of *IDH1* mutational status(15,16). Lower-grade gliomas are typically more diffuse, less proliferative, and associated with better survival compared to HGGs. Here, we use a new scalable scRNA-Seq method(17,18) for massively parallel expression profiling of human HGG surgical specimens with single cell resolution, focusing mainly on GBM. These data allow us to ask important questions such as: What is the relationship between the neural lineage resemblance of HGG cells and their proliferative status? Are transformed HGG cells directly expressing the inflammatory signatures commonly associated with certain glioma subtypes or are these expression patterns restricted to tumor-associated immune cells? Is there patient-to-patient heterogeneity in the structures of HGG cell populations? We report the broad extent of neural and non-neural lineage resemblance among transformed glioma cells, a relationship between neural lineage identity and proliferation among transformed tumor cells, and new approaches to classifying HGGs based on population structure.

## RESULTS

### Low-Cost, Scalable Single-Cell RNA-Seq of High-Grade Glioma Surgical Specimens

scRNA-Seq has emerged as a powerful approach to unbiased cellular and molecular profiling of complex tissues. Recent reports have highlighted its particular utility in solid tumors(4,15,16,19,20), where phenotypic alterations resulting from both malignant transformation and the tumor microenvironment may have great therapeutic or diagnostic significance, but are difficult to dissect from conventional bulk analysis. However, these studies employed relatively expensive and low-throughput technologies for scRNA-Seq, which complicate their sensitivity to small cellular subpopulations and ultimate routine deployment for clinical analysis. We recently reported a simple, microfluidic system for scRNA-Seq with a number of key advantages for profiling complex tissues including rapid cell loading, compatibility with live cell imaging, high-throughput (i.e. thousands of cells per sample), and low-cost without requiring cell sorting(17,18). Here, we apply this system and demonstrate routine profiling of thousands of individual cells in parallel from HGG surgical specimens. Our data set includes ~24,000 scRNA-Seq profiles from eight patients and reveals new insights into the population structures of these extremely heterogeneous tumors, relationships between neural lineages and subpopulations of transformed cells, and the immune microenvironment.

We procure tissue from surgical resections and immediately subject it to mechanical and enzymatic dissociation to produce a single-cell suspension. These cells are rapidly loaded into a microfluidic device where they are captured in arrays of microwells (**Supplementary Fig. 1A**), subjected to imaging-based quality control and automated cDNA barcoding for pooled scRNA-Seq. Importantly, we do not apply any cell sorting and attempt to randomly sample the cell suspension. **Supplementary Table 1** and **Supplementary Fig. 1B** summarize the data in terms of patient diagnosis and cell numbers, molecular, and gene detection rates, which are comparable to those obtained in previously reported, large-scale scRNA-Seq experiments in tissues(21,22), and GBM subtype as determined by comparing the single-cell average profiles to bulk RNA-Seq from previously classified GBMs in TCGA(7).

### Identification of Malignantly Transformed Glioma Cells with Single Cell RNA-Seq

We used the Phenograph implementation of Louvain community detection(23), a commonly used method for unsupervised clustering of scRNA-Seq data(24,25), to analyze the diversity of cell types in individual patients. **Figure 1A** shows t-distributed stochastic neighbor embedding (t-SNE) projections of the single-cell profiles in each patient colored based on the resulting Phenograph clusters. Differential expression analysis to identify genes specific to each cluster revealed discrete populations of endothelial cells, pericytes, T cells, myeloid cells, and oligodendrocytes (**Fig. 1A** and **Supplementary Figs. 2-9**) as expected in HGGs. In addition, each tumor harbored a large, complex population of cells comprised of multiple clusters that were often contiguous in the corresponding t-SNE projection. These cells most commonly resembled glia, expressing markers of astrocytes like *GFAP*, *AQP4*, and *ALDOC* and oligodendrocyte progenitors like *OLIG1*, *OLIG2*, and *PDGFRA*. Because transformed glioma cells typically resemble glia at the level of gene expression, we considered these cells to be putatively transformed(6) and attempted to validate these candidate transformed cells by orthogonal means. Importantly, there is no known, universal marker or set of markers that can be used for unambiguous sorting of transformed cells from glioma tissues(4). Therefore, an analytical approach for distinguishing transformed and untransformed cells, which will inevitably be mixed in our scRNA-Seq data, is crucial.

**Figure 1.**
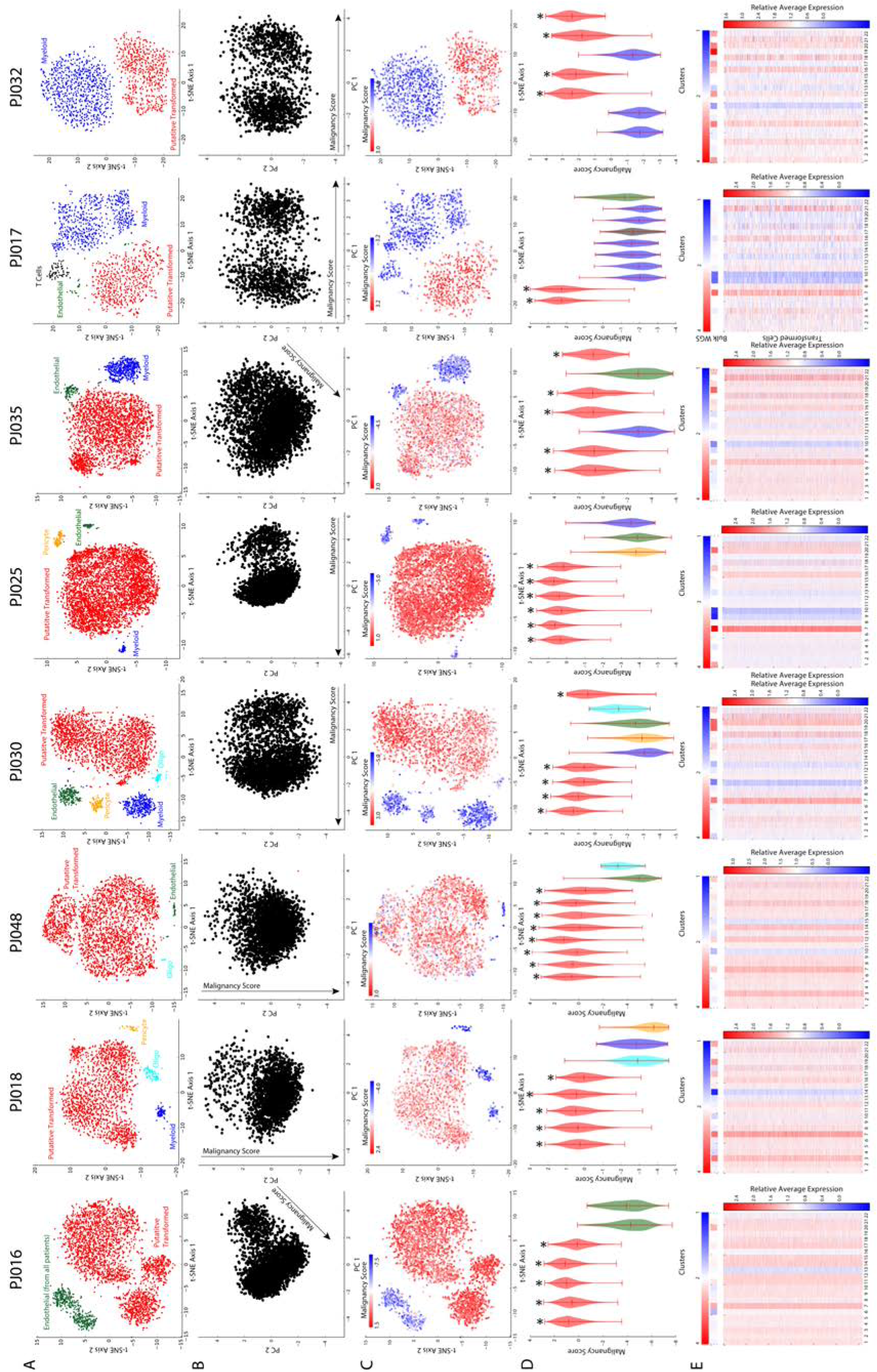
A) t-SNE projections of scRNA-Seq profiles for each tumor colored by unsupervised clustering resulting from Phenograph analysis. We note that while the putatively transformed populations in each tumor appear in red for simplicity, the majority of them actually contain multiple Phenograph clusters as shown in D) and detailed in **Figure 3**. The cell type labels are based on marker expression patterns shown in **Supplementary Figures 2-9**. B) Principal component analysis of the z-scored matrix of average chromosomal expression for each tumor showing a characteristic axis of variation, which we call the “malignancy score”, on which the putatively transformed cells are separated from the untransformed cells in each tumor. C) Same as A) but colored based on the malignancy score in B). D) Distributions of malignancy scores for each Phenograph cluster in A) showing that all of the putatively transformed clusters have higher median scores than all of the untransformed clusters within each tumor. Stars indicate the putatively transformed clusters. E) Heatmaps showing the average copy number of each chromosome based on low-pass, bulk WGS (top) and heatmaps showing the average expression of each chromosome in each cell associated with a transformed cluster relative to the average untransformed cell in each tumor (bottom).

Previous studies have shown that large copy number alterations and aneuploidies are readily detectable by scRNA-Seq of tumor tissues(4,15,16,19). We found that aneuploidies were evident in certain cellular populations based on the average expression of each chromosome in each cell. Principal component analysis (PCA) of the chromosomal expression matrix for each patient consistently revealed an axis of variation that separated the putatively transformed cells from those that expressed common markers of cells in the glioma microenvironment (**Fig 1B-C**). We called this the malignancy score, which we found to be either the first PC, second PC, or a linear combination thereof in each patient. Notably, we did not detect putatively untransformed cells in PJ016. ~90% of cells in each PJ016 cluster express *SOX2* (**Supplementary Fig. 10A**), which is normally expressed in stem and precursor cells in the CNS and, as discussed below, is pervasively expressed in transformed HGG cells compared to the adult brain. To address this, we included the combined set of endothelial cells from all of the other tumors to enable comparative analysis. Importantly, each cluster of putatively transformed cells had a significantly higher median malignancy score than the microenvironmental cells in every patient (**Fig. 1D**). Finally, when we considered the relative expression of each chromosome for each putatively malignant cell compared to the average chromosomal expression of the microenvironmental cells in each patient, we observed clear evidence of aneuploidies that are common in HGG, such as amplification of chromosome 7 and loss of chromosome 10 (**Fig. 1E**). For further validation, we conducted low-coverage whole genome sequencing (WGS) of bulk tumor tissue from each patient and computed the average copy number of each chromosome by comparison to a diploid reference. We found that the copy number variants evident in bulk WGS were in good agreement with the most prominent alterations in our scRNA-Seq data (**Fig. 1E**). As expected, there are some exceptions likely due to the lack of resolution at the single-cell level and the potential for compensatory changes in gene expression. For example, there is an apparent loss of chromosome 13 in PJ016 that is distinctly less prominent in the bulk WGS. High resolution analysis of the bulk WGS reveals that there is indeed loss of a large region of chromosome 13 (**Supplementary Fig. 11**). While this analysis is not meant to enable quantitative assessment of copy number alterations, it gives us confidence that the putatively transformed populations of cells are indeed mutated.

### SOX2 is Pervasively Expressed in High-Grade Glioma Cells

There is currently no universal marker that can consistently and specifically label transformed cells across HGGs. While it is unlikely that such a marker exists, we sought to determine whether our scRNA-Seq profiles could reveal genes that are both highly specific to and pervasively expressed in transformed glioma cells in HGG tissue. Taking advantage of the results described above, we conducted a differential expression analysis between cells in all transformed and untransformed clusters across our data set. We then asked which genes were most frequently detected in the transformed population among those with at least 8-fold higher expression (p_adj_ < 0.01) in transformed versus untransformed cells. Interestingly, we found that *SOX2*, a gene with a well-known and crucial role in stem cell biology that is commonly associated with glioma stem cells(26,27), is the most frequently detected gene with at least 8-fold specificity for the transformed cells (**Figure 2A**). Indeed, previous studies have suggested that SOX2 protein is significantly more widely expressed in glioma tissue than in normal brain, with expression in the adult brain being typically restricted to ventricular stem cell niches(28,29). We found pervasive expression of *SOX2* across all eight patients profiled in this study (**Supplementary Fig. 10**); *SOX2* is expressed by cells in every transformed cluster identified in these tumors. In fact, analysis of transcript drop-out in the transformed populations identified in these eight tumors suggests that the frequency with which we detect *SOX2* transcript likely underestimates its pervasiveness (**Figure 2B**). This is unsurprising because we have limited sensitivity, and transcription factors like SOX2 tend to be lowly expressed. To confirm these results, we carried out immunohistochemical (IHC) analysis of six of the eight tumors in our cohort and found widespread expression of SOX2 protein in every tumor (**Figure 2C**). Notably, the fraction of SOX2^+^ cells in the IHC specimens correlates strongly (r = 0.98, p = 0.001) with the fraction of transformed tumor cells inferred using our scRNA-Seq data and the analysis shown in **Figure 1** (**Supplementary Figure 12**).

**Figure 2.**
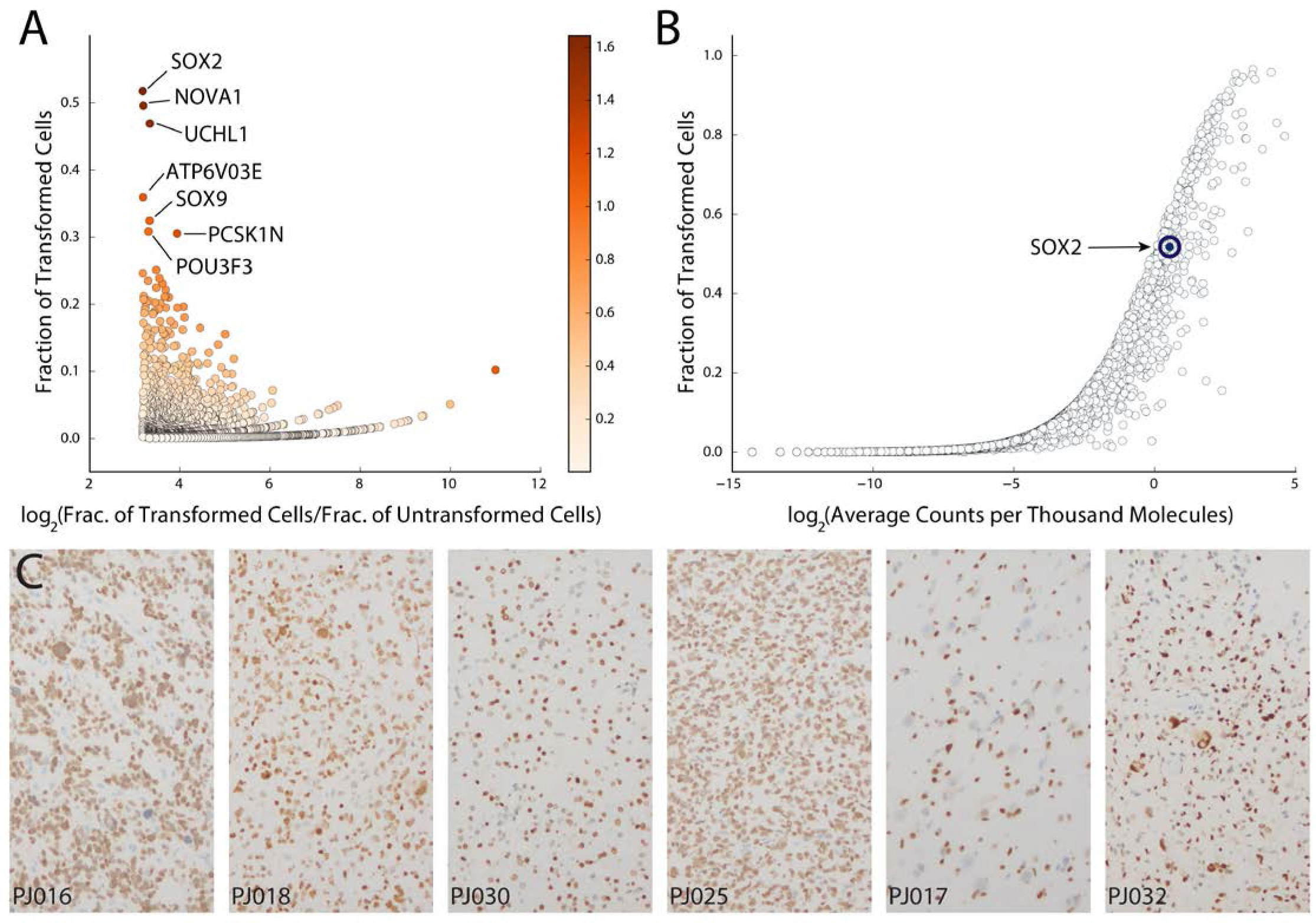
A) Analysis of the pervasiveness of genes that are highly specific to the transformed cells across all eight patients based on differential expression analysis (see Methods; all genes displayed have 8-fold specificity for the transformed cells). The colorbar represents the product of the x-and y-axes. *SOX2* is the most pervasively detected gene specific to transformed glioma cells in these eight HGG patients. B) Drop-out curve for the total population of transformed cells showing the characteristic sigmoidal shape that indicates how, for the majority of genes, higher expression (counts per thousand or CPT) leads to detection in a higher fraction of cells. Because the detection frequency of *SOX2* is close to that of similarly expressed genes, *SOX2* is unlikely to be associated with a specific subpopulation of transformed cells and the frequency with which it is expressed among transformed cells is likely to be underestimated by our data. C) IHC analysis confirming widespread protein expression of SOX2 in tissue slices from the six of the eight HGG patients in our cohort from which tissue was available for staining. We note that a considerable fraction of unstained nuclei in these specimens appear to be associated with blood vessels.

To further validate this finding, we performed quantitative immunohistochemical analysis of SOX2 expression in a larger cohort of 40 surgical specimens from high-grade gliomas (**Supplementary Figure 13**). SOX2^+^ cells were seen in all tumor samples, and the median labeling index of SOX2 in this cohort was 0.87 (**Supplementary Figure 13A**), suggesting that the vast majority of transformed HGG cells are SOX2+. SOX2 staining varied across samples, with the highest density of SOX2+ cells seen in areas of highest cellularity (**Supplementary Figure 13B**, r = 0.93, p = 10^−18^), which is also consistent with SOX2 expression in transformed HGG cells. Furthermore, regression analysis also showed a significant relationship between SOX2 labeling index and total cellularity (**Supplementary Figure 13C**, r = 0.36, p = 0.02). These results support the findings from scRNA-Seq analysis and suggest that SOX2 is a promising tumor cell marker for histopathological analysis of HGGs. In addition, we found no significant difference between primary and recurrent tumors in terms of SOX2 labeling index, suggesting that SOX2 is a useful marker of glioma cells in both pathological settings. While the pervasiveness of SOX2 is inconsistent with the idea that this protein marks a rare subpopulation of stem-like cells in HGG, the role of SOX2 in pluripotency implies that the majority of transformed glioma cells are in an immature and potentially plastic state.

### Transformed Cells Resemble both Glial and Neuronal Lineages in High-Grade Glioma

Previous studies of low-grade gliomas (LGGs) have used scRNA-Seq to draw comparisons between populations of glioma cells and certain neural lineages in the brain(15). Subpopulations resembling oligodendrocyte progenitors (OPCs) and astrocytes were reported(15). Bulk expression analysis of localized biopsies in HGG showed that different regions of the same tumor resembled disparate expression subtypes (e.g. Proneural, Classical, and Mesenchymal)(2,3) and subtype-specific differences in cellular composition and glial lineage resemblance(3). Relatedly, scRNA-Seq of relatively small numbers of cells (tens per patient) in GBM found cellular subpopulations resembling different expression subtypes cooccurring in the same tumor(4). These findings have implications for both cell-of-origin and the possibility that neurodevelopmental processes are occurring during glioma development and progression.

We sought to determine the extent to which subpopulations of transformed cells resemble neural lineages in HGGs subjected to large-scale scRNA-Seq. We performed unsupervised clustering of transformed glioma cells identified by the aneuploidy analysis described above and identified markers of the resulting subpopulations (**Figure 3A**, see Methods)(23). The heatmaps in **Figure 3B** show the expression of a subset of neural lineage markers found to associate specifically with certain subpopulations of transformed cells across our patients. As expected, we found that many of the commonly differentially expressed genes were markers of glial lineages including astrocytes (*GFAP*, *AQP4*, *ALDOC*) and OPCs (*OLIG1*, *OLIG2*, *PDGFRA*, *DLL3*). While multiple tumors contained cells that express genes associated with more mature oligodendrocytes, the tumor PJ018 contained a well-defined subpopulation that strongly resembled myelinating oligodendrocytes with specific expression of multiple myelin genes including *MBP*, *MOG*, and *MAG*. Hence, HGGs harbor cells that resemble a broad spectrum of glial developmental states and maturities. These findings are consistent with the well-established glial nature of these tumors and the numerous studies pointing to a glial cell-of-origin for gliomas(30–33). Interestingly, in multiple patients, we also observed populations of transformed cells that closely resemble immature neurons or neuroblasts - progenitors that give rise to neurons (purple boxes, **Figure 3B**). These cells express genes associated with neuroblasts like *CD24* and *STMN2* along with genes primarily expressed in the neuronal lineage in the brain, such as *DCX*(34,35). Interestingly, while we observe subpopulations that co-express these genes and have low expression of canonical glial markers (indicated by purple rectangles in **Figure 3B**), we also find subpopulations with significant co-expression of neuroblast and OPC markers (e.g. *OLIG2*, *PDGFRA*), suggesting potential plasticity between these two cell types in HGG. PJ048 harbored a particularly well-defined population of immature neuronal cells, some of which even expressed markers of more mature neurons (**Supplementary Fig. 14**). Taken together, our results indicate that the lineage resemblance of transformed glioma cells in HGG includes not only a diversity of glia, but even extends into the neuronal lineage.

**Figure 3.**
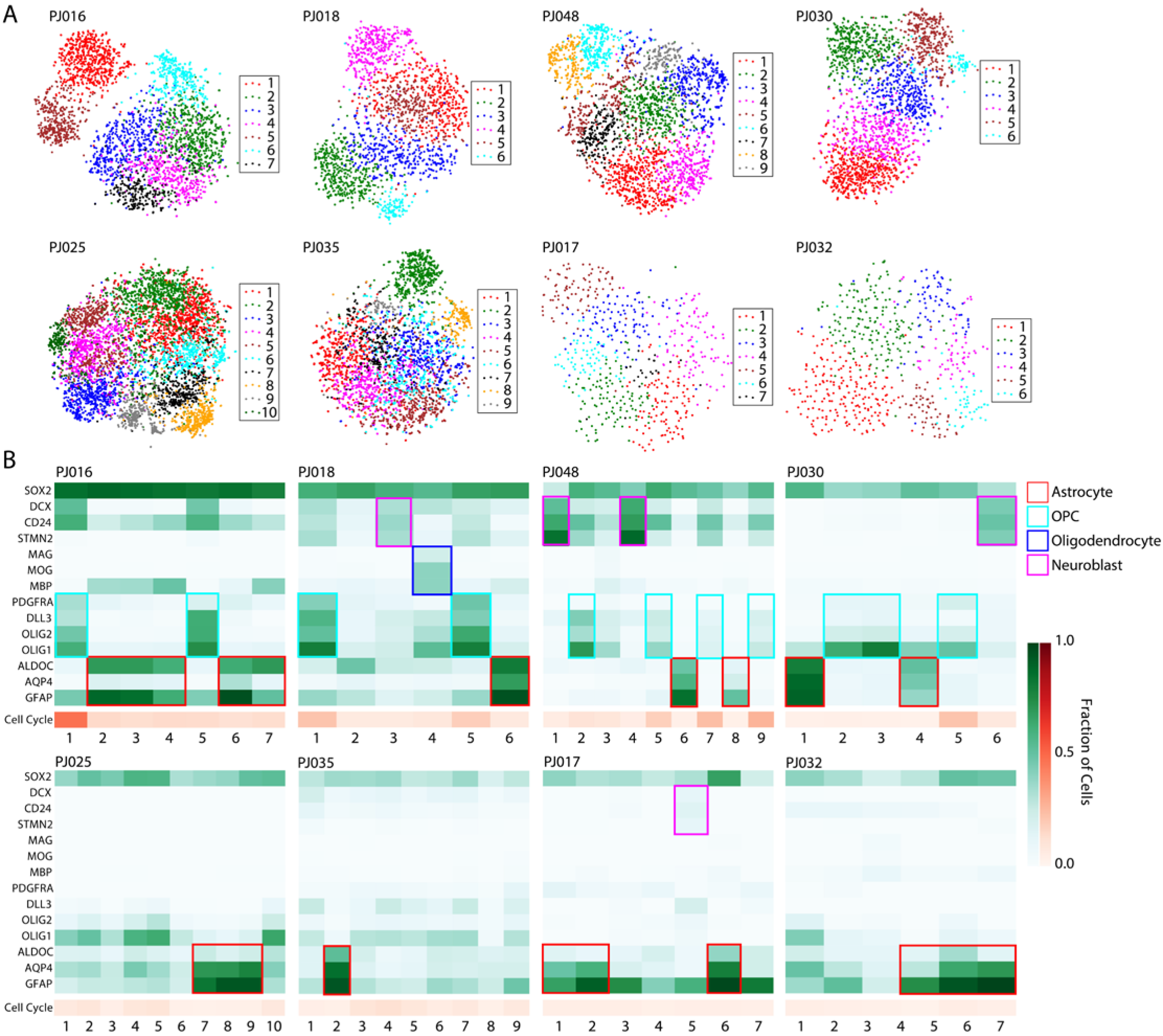
A) t-SNE projections of the transformed population of cells from each of the eight HGGs from scRNA-Seq. The projections are colored based on the cellular subpopulations identified from unsupervised clustering. B) Heatmaps showing the detection frequency of canonical astrocyte, OPC, oligodendrocyte, and neuroblast markers found to be specifically associated with transformed cellular subpopulations shown in A) across multiple patients along with *SOX2*, which is expressed across all transformed populations. The orange heatmap below each green heatmap shows the average detection frequency of cell cycle control genes found in each subpopulation. Note that some tumors have subpopulations resembling multiple neural lineages (PJ016, PJ018, PJ030, PJ048), while others exhibit a relative loss of neural lineage identity and concomitant reduction in proliferation.

### Relationship Between Proliferation and Lineage Resemblance in High-Grade Glioma

Many studies have demonstrated the efficacy of scRNA-Seq for assessing the proliferative state of individual cells and even assigning cell cycle stage(4,15,21). Recent scRNA-Seq experiments in LGGs showed that glioma cells with clear lineage resemblance to astrocytes and oligodendrocytes were generally non-proliferative, and cycling cells were largely restricted to a stem-like compartment(15). Conversely, an earlier study of GBM with scRNA-Seq reported that the stem-like glioma signature was anti-correlated with expression of cell cycle control genes, suggesting that glioma stem cells are largely quiescent in GBM(4).

We used the expression of cell cycle control genes to assess active proliferation across the subpopulations we identified in transformed cells (**Figure 3B**, **Supplementary Table 2**). Among transformed subpopulations with a clear neural lineage relationship, we never observed high expression of proliferation markers in those resembling astrocytes, myelinating oligodendrocytes, or neuroblast-like cells that lack co-expressed OPC markers. In contrast, a subset of OPC-like cells does express high levels of cell cycle control genes. These results are consistent with the behavior of these neural cell types in the adult brain, where OPCs are the predominant population of cycling cells and astrocytes, oligodendrocytes, and neuroblasts are generally not found in the cell cycle (36,37).

### Observation of Distinct Cellular Population Structures among Transformed Cells in High-Grade Glioma

Given the extent of neural lineage diversity represented in HGGs, we decided to investigate the underlying structure of the transformed population on an individual patient basis. Recent reports describe analytical methods for identifying branching events and even pseudo-temporal ordering of scRNA-Seq profiles, particularly in the context of cellular differentiation(38–41). Most of these approaches construct a graph from scRNA-Seq profiles in which each node represents a cell or group of cells and edges indicate similarity between nodes.

Our above analysis of cellular heterogeneity in HGG relies on the construction of a k-nearest neighbors graph from our scRNA-Seq profiles, which is then used for modularity clustering(23). Therefore, to visualize the relationships between subpopulations, we plotted the k-nearest neighbors graph of the transformed cells from each patient as a force-directed graph in **Figure 4** (see Methods). This analysis revealed clear differences in the structures of these populations. In particular, the transformed cells in three of the tumors formed multi-branched structures (e.g. PJ016, PJ018, and PJ048), harbored cells resembling a diversity of neural lineages, and closely resembled the Proneural subtype of GBM based on comparison of the single-cell average profiles of these tumors and classified bulk RNA-Seq data from TCGA(7). A second set of three tumors resembled the Classical subtype (PJ030, PJ025, and PJ035). PJ030 exhibited a branched structure with both OPC-and astrocyte-like branches and a small subpopulation of neuroblast-like cells, while PJ025 and PJ035 were less structured and less diverse in terms of neural lineage resemblance. Finally, PJ017 and PJ032 exhibited relatively unstructured populations and closely resembled the Mesenchymal subtype. The number of cells sampled per tumor did not explain these structural differences (**Supplementary Table 1**). In most cases, cells at the termini of the branches resemble differentiated glia. For example, PJ016, PJ018, PJ030, and PJ048 each contain a branch that terminates in a subpopulation that resembles astrocytes (**Figure 4**). PJ016 and PJ030 contain termini that resemble OPCs. The non-astrocyte branch of PJ018 strongly resembles oligodendrocyte differentiation. The terminus contains a subpopulation resembling an oligodendrocyte-like cell that expresses myelin genes and is adjacent to a subpopulation that expresses OPC markers (**Figure 4**). At the branch-point, PJ018 contains a lineage-ambiguous cell type with simultaneous expression of neuroblast and OPC markers, reminiscent of previous observations of multi-potent progenitors in the brain(42,43). PJ048 is particularly remarkable in that it harbors an astrocyte-like branch, an OPC-like branch terminating with a small population of myelin-expressing oligodendrocyte-like cells, and a large branch resembling neuroblasts or immature neurons (**Supplementary Figure 14**).

**Figure 4.**
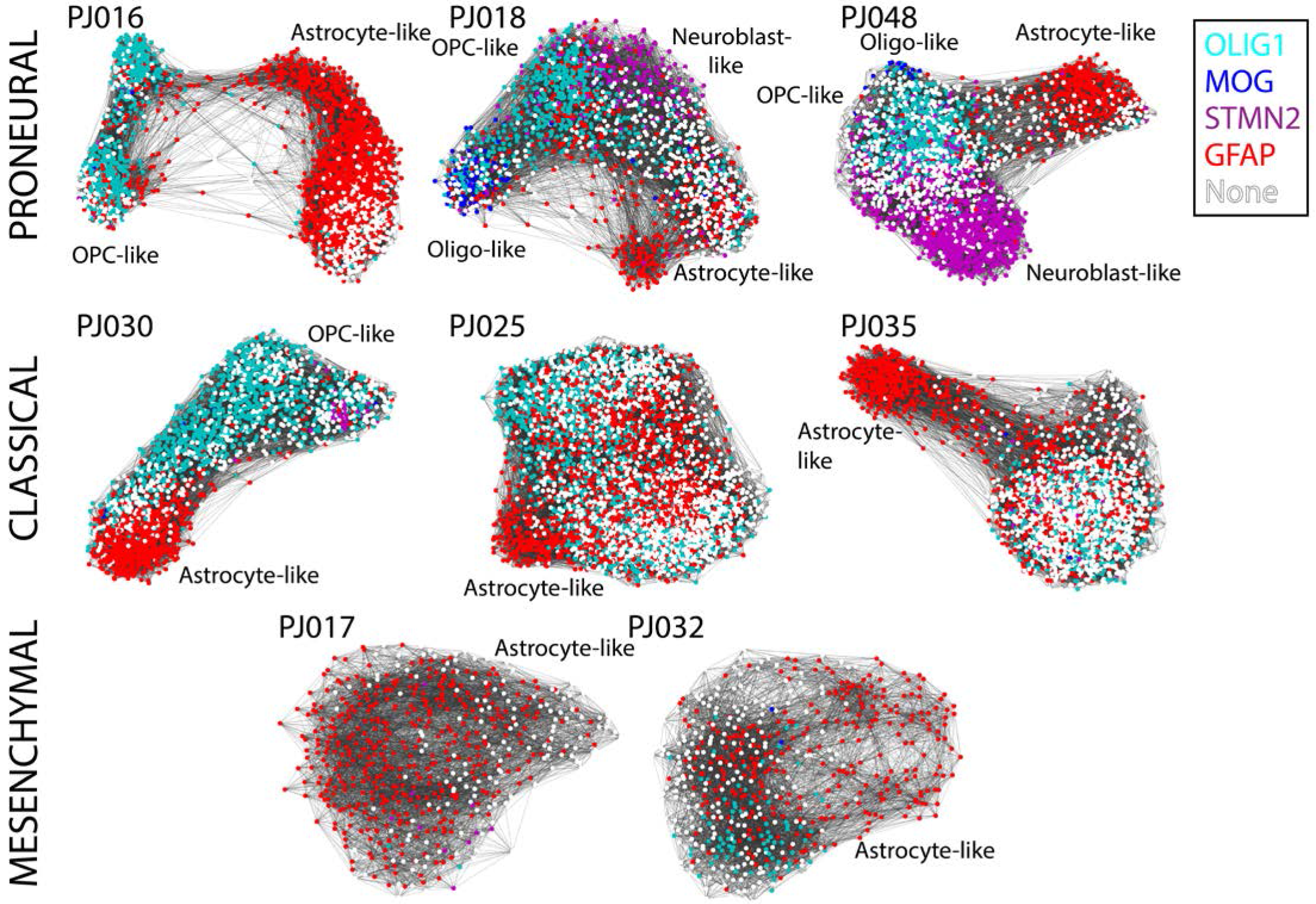
Force-directed graphs generated from the k-nearest neighbors graphs of the transformed cells profiled in each patient. Colors indicate which of the astrocyte marker *GFAP*, the OPC marker *OLIG1*, the oligodendrocyte marker *MOG*, or the neuroblast marker *STMN2* is most highly expressed in a given cell. For example, a purple cell has higher levels of *STMN2* than the other three markers. None of the four markers are detected in white-colored cells. PJ016, PJ018, and PJ048 form multi-branching structures associated specific neural lineages and their respective single-cell average profiles closely resemble the Proneural subtype of GBM. For example, one branch of PJ018 terminates with GFAP-expressing astrocytic cells, whereas the other resembles oligodendrocyte differentiation. PJ030, PJ025, and PJ035 are somewhat less structured (although PJ030 contains clearly separated OPC-and astrocyte-like branches) and have single-cell average profiles that closely resemble the Classical subtype of GBM. In contrast, PJ017 and PJ032 are unstructured, do not exhibit branching, show reduced neural lineage diversity, and have single-cell average profiles that closely resemble the Mesenchymal subtype of GBM.

While the branching behavior represents neural lineage diversity and differentiation, the cellular states of the less structured tumors are less clear. The heatmaps in **Figure 3B** show that four of the less structured tumors (PJ017, PJ025, PJ032, and PJ035) express relatively few neural lineage markers with the notable exception of astrocyte genes. We know from substantial prior work that glioma cells, and particularly GBM cells, are capable of differentiating along non-neural lineages. For example, some GBMs undergo mesenchymal transformation(44,45).

To better understand these tumors, we sought to analyze their lineage resemblance across a large database of expression profiles representing cell types in many organs. We used a curated gene expression database to identify cell types that resemble the cellular subpopulations identified among the transformed glioma cells(20). **Figure 5A** shows hierarchical clustering of correlation coefficients between the average profile of each transformed subpopulation in our dataset and the cell type-specific expression profiles in the curated database. This analysis immediately reveals three clusters of cell type-specific expression profiles - one enriched in embryonic stem and neural cells (neural/ESC), one enriched in immune cells (immune), and one enriched in mesenchymal and mesenchymal stem cells (mesenchymal/MSC) – along with three multitumor groups of transformed subpopulations. The first group of transformed cells (Group I) is exclusively comprised of subpopulations from branched tumors PJ016, PJ030, and PJ048. Glioma cells in Group I are correlated with the ESC/neural cluster, but bear the weakest resemblance to the MSC/mesenchymal and immune clusters. Group II contains clusters from all but one tumor (PJ032, a recurrent GBM) and is correlated with the neural/ESC signature, but unlike Group I has some mesenchymal/MSC and immune character. Interestingly, Group III is comprised of subpopulations from only two tumors, PJ017 and PJ032, which strongly resemble both the immune and MSC/mesenchymal clusters, but weakly resemble the ESC/neural cells.

**Figure 5.**
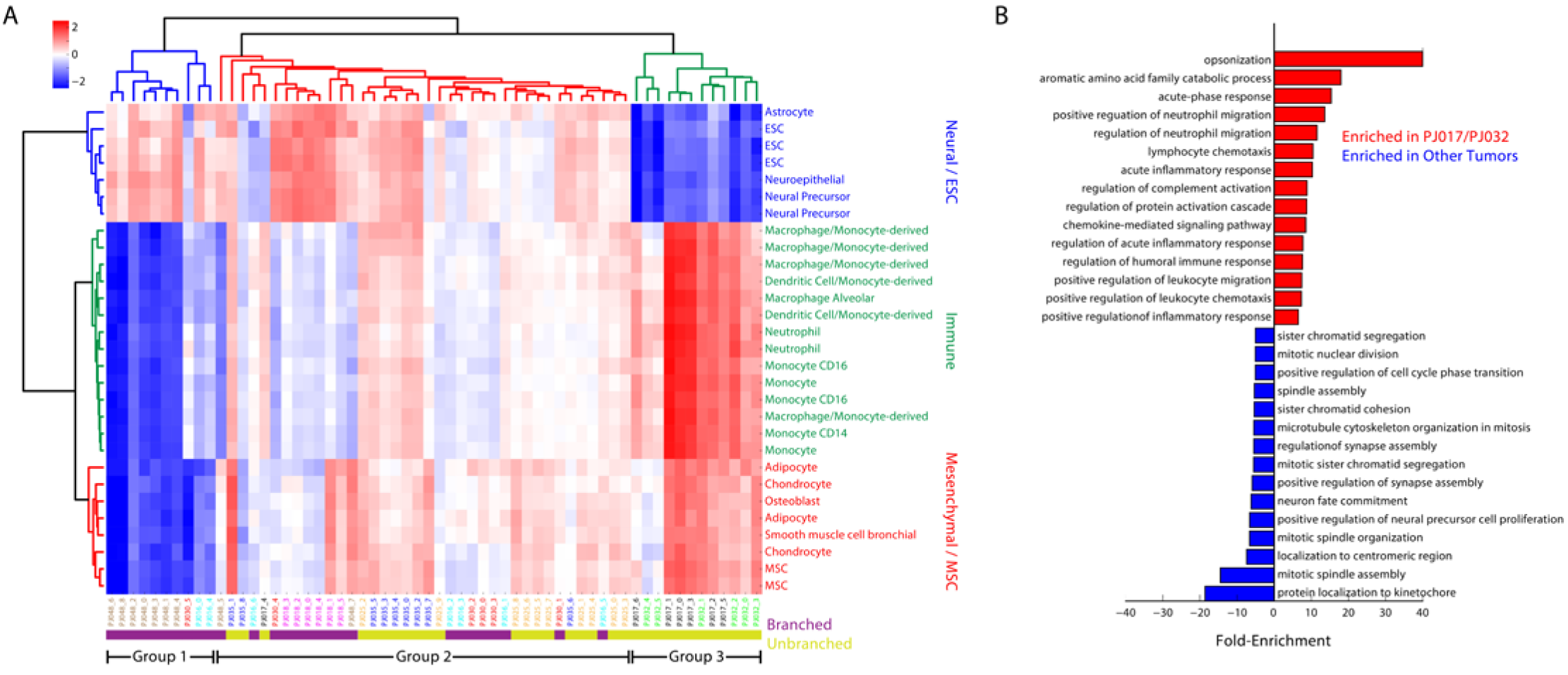
A) Hierarchical clustering of the correlation between each transformed subpopulation and a database of cell type-specific expression profiles with high variability across the data set. We find three cell type clusters referred to as Neural/ESC, Immune, and Mesenchymal/MSC which divide the tumor cell subpopulations into three major groups. B) Gene ontology analysis of the differentially expressed genes between the Group III tumors (PJ017/PJ032) and the remaining tumors (PJ016, PJ018, PJ025, PJ030, PJ035, PJ048) after removal of genes specific to the untransformed immune cells in PJ017 and PJ032. The Group III tumors show a clear immunological gene signature that is specific to the transformed cells.

The results in **Figure 5A** are consistent with the notion that the tumors lacking clear neural lineage structure have undergone mesenchymal transformation. However, this analysis also highlights crucial distinctions among these tumors. First, as has been recognized from bulk expression analysis of GBM, mesenchymal gene expression is often accompanied by expression of inflammatory genes(3,7). However, the extent to which this inflammatory signature is expressed by transformed glioma cells has been difficult to discern from bulk analysis due to the presence of both transformed and untransformed cells. Here, we find that mesenchymally transformed glioma cells express many immune-related genes, but that there is also significant variability in the expression of these genes among subpopulations with mesenchymal gene expression (Group II vs. Group III). Second, the two tumors with high levels of inflammatory gene expression (PJ017 and PJ032 in Group III) also bear the least resemblance to the neural/ESC clusters. Hence, expression of this mesenchymal-associated immune signature is accompanied by loss of neural lineage identity.

PJ017 and PJ032 are notable not just because of their unbranched structure, strong immunological gene expression, and loss of neural lineage identity, but also because they are the only two tumors in our data set where transformed glioma cells are in the minority of profiled cells. PJ017 is 48% myeloid cells, 5% T cells, and 45% transformed glioma cells; PJ032 is 57% myeloid cell and 43% transformed glioma cells based on scRNA-Seq. While the observation of extensive myeloid infiltration in tumors that express high levels of inflammatory markers is intriguing, it also raises the possibility that our observation arises from experimental cross-contamination either during mRNA capture or library construction. We compared the transformed cells in PJ017/PJ032 to the remaining tumors after stringent filtration of the differentially expressed genes to remove any genes that are more highly expressed in the myeloid compartment of these tumors and could result in cross-contamination (see Methods). **Figure 5B** shows that, despite our stringent filter, the transformed cells in PJ017/PJ032 express high levels of immune genes compared to the remaining tumors (**Supplementary Table 3**), thus indicating that tumor cells expressing an immune-like signature may recruit infiltration of myeloid cells. Interestingly, we were able to validate this finding in an independent patient cohort (**Supplementary Figure 15**) by re-analyzing an earlier, smaller-scale GBM data set from Patel *et al* where one out of the five tumors profiled expressed this same signature at high levels among transformed cells(4).

We next asked if any of the genes in this signature have known receptor-ligand interactions with cognates expressed in the myeloid cells of these tumors. One interaction of potential therapeutic interest in glioma is the macrophage proliferation cytokine CSF1 and its cognate receptor CSF1R, which has been extensively validated in pre-clinical studies in glioma along with its potential therapeutic efficacy(46,47). **Figure 6** shows that *CSF1R* is widely expressed in the myeloid populations in the seven tumors in which we detect myeloid cells. However, *CSF1* is most highly expressed in the transformed glioma cells PJ017 and PJ032, the two tumors with the highest immune signature correlation in **Figure 5A** and the highest proportion of tumor-associated myeloid cells. These results are consistent with a model in which CSF1 secretion by glioma cells recruits CSF1R-expressing microglia or macrophages to the tumor microenvironment, as demonstrated previously in murine models(46,47), and may point to a patient population that would be particularly susceptible to CSF1R blockade.

**Figure 6.**
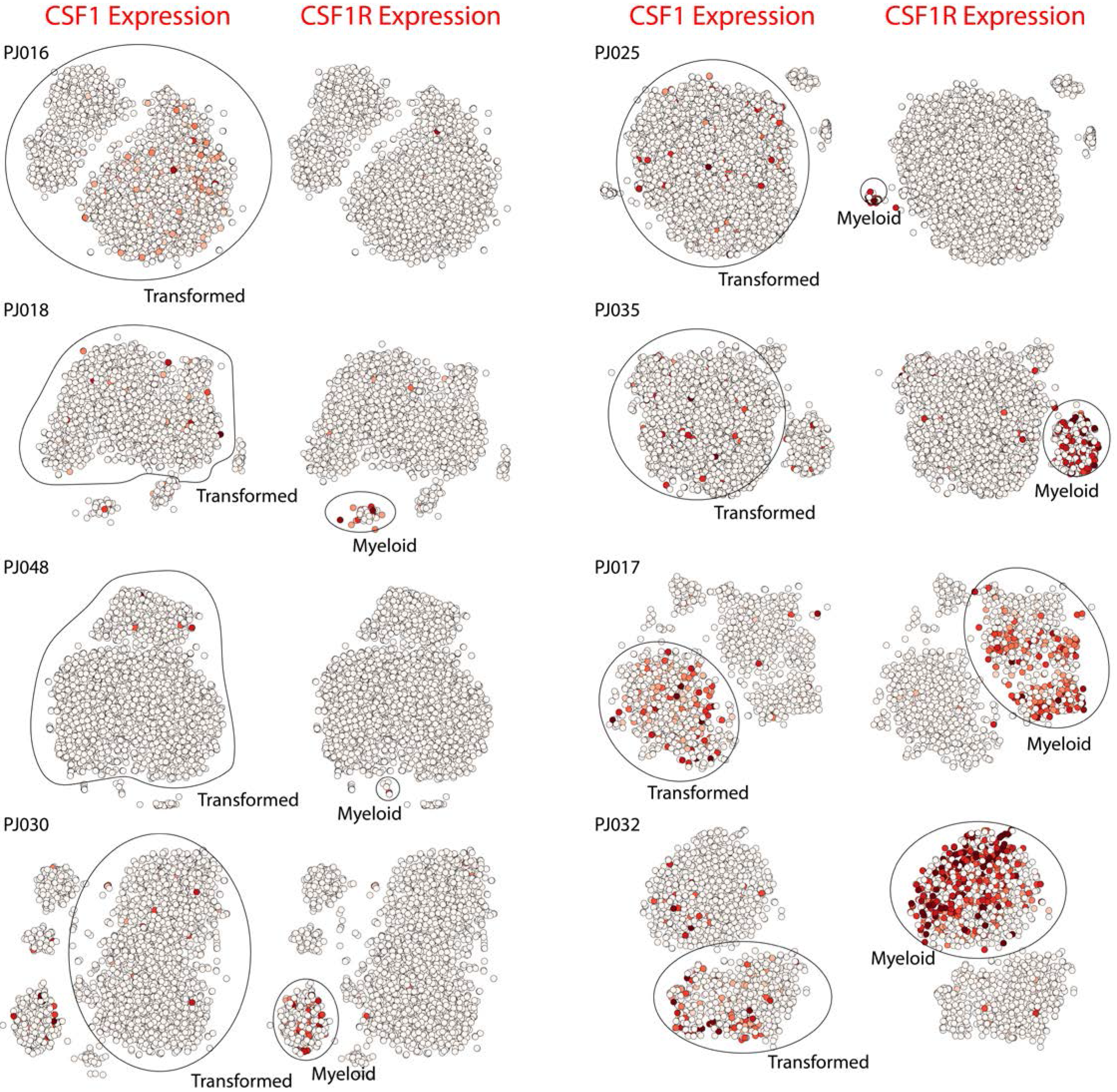
t-SNE projections of scRNA-Seq profiles from all eight tumors. The plots are colored by expression of either *CSF1*, a macrophage stimulating cytokine, or the gene encoding its cognate receptor *CSF1R*. Receptor expression is widespread among myeloid cells, but expression of the cytokine is significantly higher in the transformed glioma cells of PJ017 and PJ032 than in the other tumors. We note that no myeloid cells were detected in PJ016.

Previous studies have used scRNA-Seq to analyze the heterogeneity of tumor-associated myeloid cells in gliomas. One study focusing on *IDH1* mutant gliomas found a continuous distribution of myeloid phenotypes ranging from more microglial on one extreme to more macrophage-like on the other(16). In contrast, a recent study focusing on HGGs found a clear separation between cell of microglial origin and blood-derived macrophages(48). We applied similar analytical methods in our patient cohort and found significant differences in myeloid phenotype dominated mainly by expression of pro-inflammatory cytokines and microglial versus macrophage lineage resemblance (**Supplementary Figure 16-18**).

## DISCUSSION

Large-scale scRNA-Seq has allowed us to dissect the lineage identity and proliferative status of malignant cells in HGG with unprecedented resolution. We find that only a subset of transformed cells resembles neural lineages, and that there is significant inter-tumoral heterogeneity in the diversity of neural lineages represented among transformed cells. Furthermore, we find that neural lineage resemblance extends beyond glia. Subpopulations of transformed cells in multiple patients resemble neuroblasts or immature neurons. Moreover, we defined a molecular classification for HGGs based on population structure at the single cell level that is closely related to the range of neural lineage resemblance among transformed cells in a tumor. Specifically, transformed populations with branched structures resemble a variety of neural lineages arranged similarly to normal neurodevelopment. These transformed populations appear to obey the same rules for proliferative potential as their corresponding neural lineages in the brain. Based on expression of cell cycle genes, the transformed cells resembling astrocytes, myelinating oligodendrocytes, and neuroblasts are generally not in the cell cycle, whereas those resembling OPCs are often found in a proliferative state. These observations are distinct from what has been reported previously in low-grade oligodendrogliomas, where a truncal, stem-like population was found to encompass the cycling cells in a tumor(15). In HGG, we find that proliferative state is predominantly associated with cells expressing markers of OPC-like, glial progenitors.

A second group of tumors harbored transformed cells that either resemble astrocytes or exhibit a loss of neural lineage identity. The underlying subpopulations tend to resemble mesenchymal and immune cell types, and express low levels of proliferation markers. However, there is significant inter-tumoral heterogeneity among these HGGs, particularly with respect to immunological gene expression and corresponding myeloid infiltration. Mesenchymal gene expression in HGGs has long been associated with an inflammatory gene signature based on bulk analysis of tumor tissue(3,7). Indeed, previous studies have shown the essential role of inflammation-associated transcription factors such as STAT3 and CEBPB in mesenchymal transformation(45). Here, we define a mesenchymal-associated immunological signature expressed specifically by transformed glioma cells in a subset of patients. In addition, we find that these tumors express high levels of the macrophage recruitment factor gene *CSF1*, a cytokine whose cognate receptor *CSF1R* is widely expressed in myeloid cells across our patient cohort. One intriguing possibility is that CSF1 secretion by HGG cells is responsible for enhanced myeloid infiltration and that CSF1R blockade, which has been investigated as potential therapy for HGG(46,47), would be particularly beneficial to this subset of patients.

## CONCLUSION

The combined insights into both transformed population structure and microenvironment, even in the context of a modest cohort and a disease with extensive molecular characterization, highlight the utility of large-scale scRNA-Seq in complex tumors. We anticipate that the rapid, scalable, and inexpensive assessment of cellular composition, proliferative potential, tumor cell phenotype, and expression of therapeutic targets afforded by this approach will play a crucial role in molecular diagnosis and precision oncology for HGGs.

## METHODS

### Procurement and Dissociation of High-Grade Glioma Tissue

Single-cell suspensions were obtained using excess material collected for clinical purposes from de-identified brain tumor specimens. Donors (patients diagnosed with HGG) were anonymous. Tissues were mechanically dissociated to single cells following a 30 minute treatment with papain at 37C in Hank’s balanced salt solution. After centrifugation at 100 × g, the cell pellet was re-suspended in Tris-buffered saline (TBS, pH 7.4) and red blood cells were lysed using ammonium chloride for 15 minutes at room temperature. Cells were washed in TBS, counted, and re-suspended in TBS at a concentration of 1 million cells per milliliter for immediate processing.

### Massively Parallel Single-Cell RNA-Seq

We used a previously reported, automated microwell array-based platform for pooled scRNA-Seq library construction and followed the procedures for device operation, library construction, and sequencing described by Yuan and Sims(18) with the following two modifications: 1) Live staining of single cell suspensions was performed on ice for 15-30 minutes; 2) aliquots of amplified cDNA were pooled together before purification with Ampure XP beads (Beckman). Each device contained 150,000 microwells (50 μm diameter, 58 μm height) with center-to-center distance of 75 μm.

### Low-Pass Whole Genome Sequencing

For each tumor, a 2-3 mm^3^ piece was used for DNA extraction. Each section was re-suspended in 400 μL of DNA/RNA Lysis Buffer (Zymo) and homogenized with a Dounce homogenizer if necessary. DNA and RNA were then extracted for the tissue using the ZR-Duet Kit (Zymo) according to the manufacturer’s instructions. DNA was quantified using the Qubit dsDNA High Sensitivity Kit (ThermoFisher Scientific). Libraries for low-pass WGS were constructed using by *in vitro* transposition using the Nextera XT kit (Illumina). DNA inputs for each sample were normalized to 1 ng and library preparation was performed according to the manufacturer’s instructions, using unique i7 indices for each sample. Libraries from all eight tumors were pooled at equimolar concentrations, denatured, diluted, and sequenced on an Illumina NextSeq 500 using a 150 cycle High Output Kit (Illumina, 2×75 bp).

### Whole Genome Sequencing Analysis

Reads were aligned to the human genome (hg19) using bwa-mem, and coverage at each nucleotide position was quantified using bedtools after removing PCR duplicates with samtools. To generate the bulk WGS heatmaps in **Figure 1E**, we computed the number of de-duplicated reads that aligned to each chromosome for each piece of tumor tissue and divided this by the number of de-duplicated reads that aligned to each chromosome for a diploid germline sample from one of the patients (pooled blood mononuclear cells). We then normalized this ratio by the median ratio across all chromosomes to estimate the average copy number of each chromosome.

### Immunohistochemical Analysis

Immunohistochemistry using standard immunoperoxidase staining was performed on formalin-fixed paraffin-embedded tissue sections (5 microns thick) from specimens of each of the tumor resections. Briefly, we used 3×3 min cycles of de-paraffinization in xylene, 2×1 min cycles of dehydration in 100% ethanol, 2×1 min cycles of dehydration in 95% ethanol, and a one minute cycle of dehydration in 70% ethanol. Slides were then washed in water. We used 0.01M citrate buffer (pH 6) for antigen retrieval in a microwaved pressure cooker for 20 minutes. We then washed the slides three times in phosphage-buffered saline (PBS) after cooling for 30 minutes. We quenched endogenous peroxidase in 3% hydrogen peroxide in PBS for 10 minutes, washed three times in PBS, and blocked with 10% goat serum for 25 minutes. We then incubated the slides with primary antibodies for 90 minutes at room temperature. We used the following primary antibodies: rabbit anti-CD163 (Abcam, ab182422, 1:50 dilution), rabbit anti-SOX2 (Abcam, ab92494, 1:100 dilution), rabbit anti-TMEM119 (Abcam, ab185333, 1:300 dilution). After washing three times in PBS, we incubated the slides with biotinylated goat anti-rabbit secondary antibody (Vector Laboratories, 1:200 dilution) for 30 minutes at room temperature, followed by additional PBS washing, 30-minute incubation with ABC peroxidase reagent, development in DAB-peroxidase substrate solution (DAKO), and counter-staining in hematoxylin.

For the SOX2 validation cohort, tissue samples from 40 surgical resections of HGG (29 primary and 11 recurrent tumors) were fixed in 10% formalin and embedded in paraffin for immunohistochemical analysis. Five-micrometer sections were immunostained for SOX2 and counter-stained with hematoxylin. The slides were then scanned and digitized at 40x magnification on a Leica SCN400 system (Leica Biosystems). Total cell density and SOX2+ nuclei were measured using a semi-automated cell-counting algorithm as previously described(49). Algorithm-derived cell counts were manually verified, and total cell density and SOX2 cell density were assessed for one representative high-power field from each sample. The labeling index was computed by dividing the total number of SOX2^+^ cells by the total cell count for each high-power field.

### Single Cell RNA-Seq Alignment and Data Processing

As previously described, cell and molecular barcodes are contained in read 1 of our paired-end sequencing data, while all genomic information is contained in read 2(18). We trimmed read 2 to remove 3’ polyA tails (>7 A’s), and discarded fragments with fewer than 24 remaining nucleotides. Trimmed reads were aligned to GRCh38 (GENCODE v.24) using STAR v.2.5.0 with parameters “`–sjdbOverhang 65 –twopassMode Basic –outSAMtype BAM Unsorted`”(50). Only reads with unique, strand-specific alignments to exons were kept for further analysis.

We extracted 12-nt cell barcodes (CBs) and 8-nt unique molecular identifies (UMIs) from read 1. Degenerate CBs containing either ‘N’s or more than four consecutive ‘G’s were discarded. Synthesis errors, which can result in truncated 11-nt barcodes, were corrected similarly to a previously reported method(24). Briefly, we identified all CBs with at least twenty apparent molecules and for which greater than 90% of UMI-terminal nucleotides were ‘T’. These putative truncated CBs were corrected by removing their last nucleotide. This twelfth nucleotide became the new first nucleotide of corresponding UMIs, which were also trimmed of their last (‘T’) base.

All reads with the same CB, UMI, and gene mapping were collapsed to represent a single molecule. To correct for sequencing errors in UMI’s, we further collapsed UMI’s that were within Hamming distance one of another UMI with the same barcode and gene. To correct for sequencing errors in cell barcodes, we then collapsed CBs that were within Hamming distance one of another barcode, had at least 20 unique UMI-gene pairs, and had at least 75% overlap of their UMI-gene pairs. Finally, we repeated UMI error correction and collapse using the error-corrected CBs. The remaining barcode-UMI-gene triplets were used to generate a digital gene expression matrix.

### Filtering Cell Barcodes

We estimated the number of cell barcodes corresponding beads associated with cells in our microwell system using the cumulative histogram of reads associated with each barcode as described previously(21). To avoid dead cells and library construction artifacts, we removed cell barcodes that failed to satisfy certain criteria. We removed all cells where >10% of molecules aligned to genes expressed from the mitochondrial genome or where the ratio of molecules aligning to whole gene bodies (including introns) to molecules aligning exclusively to exons was >1.5. These measures remove cells with compromised plasma membranes, which results in retention of mitochondrial or nuclear transcripts(51). We also removed cells where the number of reads per molecule (indicative of amplification efficiency) or the number of molecules per gene deviated by more than 2.5 standard deviations from the mean for a given sample.

### Unsupervised Clustering, Differential Expression, and Force-Directed Graphical Analysis

To calculate k-nearest neighbors graphs, we computed a cell by cell Spearman’s correlation matrix for each population and set k=20. Spearman’s correlation was calculated from a set of genes selected because they were detected in fewer cells than expected given their apparent expression level. For this step of our analysis only, molecular counts within each column of gene by cell expression matrices were normalized to sum to 1. Genes were then ordered by their mean normalized value in the population and placed into bins of 50 genes. A gene’s detection frequency was calculated as fraction of cells in which at least one molecule of a gene was detected, and its score was defined as the maximum detection frequency in its bin minus its detection frequency. Genes with scores greater than 0.15 were considered markers and used to compute Spearman’s correlation.

This k-nearest neighbors graph was used as input to Phenograph(23), a modularity-based clustering algorithm. The similarity matrix described above was converted to a distance matrix, and used as input to tSNE(52) for visualization. Differential expression analysis was conducted using a binomial test as previously described(24).

Force-directed graphs were generated from the k-nearest neighbors graphs described above using the *from_numpy_matrix*, *draw_networkx*, and *spring_layout* commands in the NetworkX v1.11 module for Python with default parameters.

### Identification of Transformed Cells by Single Cell Analysis of Copy Number Alterations

For unsupervised identification of transformed cells in our HGG data, we first converted the raw molecular counts for each cell to log2(counts per thousand molecules + 1). We then discarded all genes that were expressed in fewer than 100 cells per tumor as well as the HLA genes on chromosome 6, which could manifest as copy number variants particularly in myeloid populations. Next, we computed the average of log_2_(counts per thousand molecules + 1) across the genes on each somatic chromosome, resulting in an N×22 matrix, where N is the number of cells. Finally, we z-scored the resulting profile for each cell and computed the principal components (PCs) of the resulting z-matrix. For each tumor, either the first PC (PJ017, PJ025, PJ030, PJ032), second PC (PJ018, PJ048), or the sum of the first two PCs (PJ016, PJ035) yielded an axis along which the putatively transformed and untransformed cells identified by clustering were separated (**Figure 1B**) as evidenced by the t-SNE projections in **Figure 1C** in which the cells are colored based on their value along the appropriate axis. To compute the heatmaps of chromosomal gene expression in **Figure 1E**, we took the average value of log_2_(counts per thousand molecules + 1) for each chromosome in each transformed cell and divided by the average value of log_2_(counts per thousand molecules + 1) for each chromosome averaged over all untransformed cells in a given tumor.

### Subpopulation Clustering with Reference Component Analysis Database

To identify cell types resembling the transformed subpopulations that we identified across our data set, we used the RNA-Seq databases curated for Reference Component Analysis(20). We first removed all transcriptomes of whole homogenized tissues (e.g. whole brain, whole blood, etc.) or that originated from cancers. We then computed the Spearman’s correlation coefficient between each cell type-specific transcriptome and the average expression profile of each transformed subpopulation across all eight tumors in our data set. All cell types with a below-median standard deviation were then removed to enrich for cell types with high variation across our data set, and the resulting correlation matrix was standardized and subjected to hierarchical clustering with a Euclidean distance metric using the *clustermap* function in the Seaborn Python module (**Figure 5A**).

We have previously shown that molecular cross-contamination in our microfluidic system is ~1%. Such a cross-contamination rate slightly reduces the contrast in gene expression profiles between different cell subpopulations. However, it is unlikely that a gene would become highly differentially expressed in, and hence become a marker of, a population of cells due to cross-contamination. To address the possibility that the immune signature that is highly enriched in PJ017 and PJ032 arises from cross-contamination due to the high abundance of myeloid cells in these two tumors, we conducted an orthogonal and more direct analysis to determine whether or not glioma cells in PJ017 and PJ032 express higher levels of immune genes than other tumors. We first conducted a differential expression analysis between the combined transformed cells from PJ017/PJ032 and the remaining tumors along with parallel analyses comparing the transformed cells from PJ017 or PJ032 to their respective immune populations using the binomial test described above. We then selected all of the genes that were significantly more frequently detected in the PJ017/PJ032 transformed cells (p<0.01 with fold-enrichment > 10) than other tumors and removed all genes that were more frequently detected in the immune cells in either tumor. Any remaining differentially expressed genes are more highly expressed in the transformed cells from PJ017/PJ032 and therefore cannot arise from molecular cross-contamination. Finally, we conducted a gene ontology analysis on the remaining differentially expressed genes that were either more frequently detected in PJ017/PJ032 (after filtering immune cell-specific genes) or in other tumors using Panther (www.pantherdb.org). **Figure 5B** shows the results for the lowest-level gene ontologies (top 15 biological process ontologies for each group) based on the Panther gene ontology hierarchy (to avoid the use of extremely broad ontologies like “cell part”, etc.).

### Generation of Myeloid Signatures

To generate microglial-and macrophage-specific gene signatures for **Supplementary Figure 17H**, we started with the cell type-specific gene sets obtained from murine lineage-tracing studies (Supplementary Table S4 from Bowman et al)(14), similar to previous analysis(48). We then assembled all of the myeloid and non-myeloid cells in our data set and conducted a differential expression analysis using the binomial test described above to identify genes with at least 5-fold specificity for the myeloid population and FDR < 0.01. We removed any genes from the Bowman et al gene sets that did not intersect with this list to avoid inclusion of genes expressed in other cell types (particular the transformed cells) to obtain the gene sets in **Supplementary Table 4**.

## DECLARATIONS

### Ethics Approval and Consent to Participate

Tissue for these studies was procured from de-identified patients through a protocol approved by the Columbia Institutional Review Board (IRB).

### Availability of Data and Materials

Raw and processed data will be made publicly available on the Gene Expression Omnibus under accession GSE103224.

### Author Contributions

JY conducted the scRNA-Seq experiments. JNB, JS, and PC procured the tissue. AL, VF prepared the tissue for scRNA-Seq. JY, HML, and PAS conducted the computational data processing and analysis with significant assistance and input from MC, AL, and AI. ECB conducted the DNA sequencing experiments. AD, GZ, DMB, and PC conducted the IHC analysis. All authors contributed to writing the paper.

## Acknowledgements

The authors thank the Molecular Pathology Core of the Herbert Irving Comprehensive Cancer Center and the Sulzberger Columbia Genome Center for technical assistance with immunohistochemistry and sequencing, respectively.

